# Artificial introns for effective expression of transgenes in mammalian cells

**DOI:** 10.1101/2021.09.01.457939

**Authors:** D.S. Naberezhnov, E.A. Lesovaya, K.I. Kirsanov, M.G. Yakubovskaya

## Abstract

Introns are widely used in the assembly of genetic constructions expressing transgenic proteins in eukaryotic cells for the enhancement of this expression. However, the choice of introns that can be applied for such purposes is limited by the excessively large size of the majority of natural introns (several thousand nucleotides) and therefore they cannot be cloned in a genetic construction. With the help of site-directed mutagenesis we have generated a library of short (99 nucleotides long) introns. The efficiency of these introns in the enhancement of gene expression was analyzed. As a result, a set of 12 introns was selected. The generated intros can be used for genetic constructions with high expression level of recombinant proteins.

## Introduction

Introns can significantly enhance the level of gene expression. For the first time this phenomenon was found in maize [1] and later was named intron-mediated enhancement of gene expression (IME) [2]. Afterwards this effect was detected in all eukaryotes [3–5]. The mechanism of IME does not depend on the splicing efficiency and is not yet fully understood [6,7]. Some investigations show that IME can be associated with the interaction of the small nucleoprotein U1 enclosed in the structure of small nucleoprotein spliceosome with 5’ splice site (5’-SS) [8–10], other splicing-involved proteins [11], histone acetylation [12], formation of DNA loops [13], mRNA export from the cell nucleus [14,15], influence on translation through exon junction complex (EJC) [16].

Due to the requirement for high level of transgene expression in the molecular biological and biotechnological research, IME became widespread as a tool for expression enhancement. Small t-intron, SV40 intron, rabbit β-globin intron, chimeric intron, and hybrid intron were employed for these purposes [17]. These introns represented full-size or partly truncated by excision of the internal part natural introns.

By means of rational design and site-directed mutagenesis we obtained a set of new artificial introns not existing in nature, but containing elements universal for all introns.

Elements essential for intron splicing differ depending on intron type; for introns of protein coding genes the essential elements are represented by donor and acceptor splice sites 5’-SS and 3’-SS, branching point, and polypyrimidine sequence [18,19] (Fig. 1).

**Figue 1.**
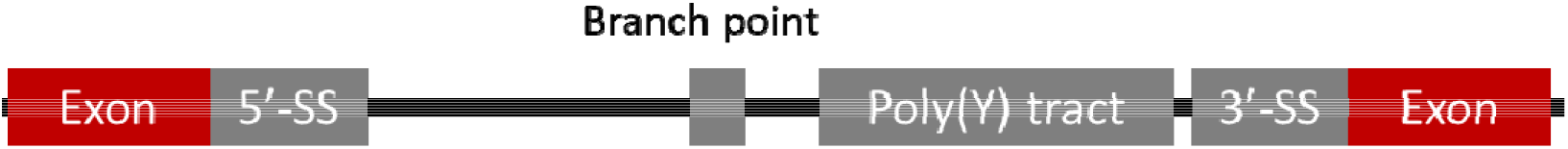
General scheme of intron.

In the 5’-SS of intron the sequence of nucleoprotein U1 RNA binding is localized [20], 3’-SS sequence and polypyrimidine sequence contain sequences of small (35 kDa) subunit U2AF binding [21] and polypyrimidine tract-binding protein (PTB) responsible for splicing regulation binding [22] respectively, and the branching point contains the sequence of large (65 kDa) subunit U2AF binding [23]. Artificial introns must contain all these elements for effective splicing.

Natural introns are characterized by wide distribution of lengths from 50 to several thousand nucleotides with the pick value of 100-200 nucleotides [24] and diverse splicing efficiency correlated with the length and IME [25,26]: longer introns demonstrate better efficiency. For molecular biological investigations short introns are preferable, therefore we have chosen the length of 99 nucleotides for the generation of artificial introns.

## Materials and methods

### Molecular cloning

Plasmids were obtained by standard restriction – ligation method and Golden Gate method. PCR conditions, sequences of plasmids and primers are presented in the Supplement.

The library of plasmids pSB/IR-CA-HybIntr-2×99intr-SV40intr-BleoR-T2A-mCherry was obtained by cloning of the library of DNA duplexes generated by renaturation of oligonucleotides 130-99intr-F and 131-99intr-R (with completion of complementary DNA T4 strands with DNA-polymerase and restriction by endonucleases BsaI (NEB, USA) and KpnI) into pSB/IR-CA-HybIntr-SV40intr-BleoR-T2A-mCherry plasmid restricted with restriction endonucleases BsaI (NEB, USA) and AsiG.

#### Cell line culturing

HEK293T cell line was cultivated in 24-well plates in DMEM medium (Paneco, Russia) with 10% calf embryonic serum (Paneco, Russia) in the incubator with 5% CO_2_ at 37° C.

#### Transfection

5 hours before transfection the cell culture medium was changed. The cells were transfected with the help of TurboFect (Thermo Fisher Scientific, USA) according to the manufacturer’s instructions. Plasmid DNA for transfection was isolated using Plasmid Miniprep Kit (Evrogen, Russia) according to the manufacturer’s instructions. In 2 days the medium was changed, and a new portion with 100 mcg/ml antibiotic zeocin was added.

#### Fluorometry

Fluorometry was performed on Fluoroskan FL (Thermo Fisher Scientific, USA) with FL1 filters.

## Results

Artificial introns (99 intron N, where N is the serial number of intron) were generated by site-directed mutagenesis with subsequent selection of variants capable of undergoing splicing. For this purpose, we designed a plasmid pSB/IR-CA-HybIntr-SV40intr-BleoR-T2A-mCherry in which under the control of promoter CAG the gene of zeocin resistance fused through T2A peptide with mCherry protein was placed (Fig. 2). Such plasmid construction is designed so that the artificial intron insertion splits the reading frame of zeocin resistance gene, and if the intron is not capable of splicing the resistance gene remains inactive. This plasmid construction allows selection of intron variants capable of splicing with the help of antibiotic zeocin.

**Figure 2.**
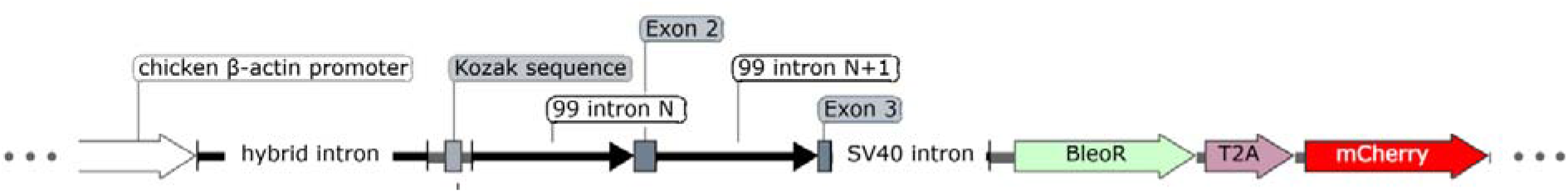
Scheme of expression cassette applied for the selection of artificial introns.

The general intron design was developed based on acknowledged literature data. GTAAGT sequence complementary to the sequence of spliceosome RNA U1 was chosen as 5’-SS intron sequence [27]. CAG sequence common for all introns was used as 3’-SS, and TACTAAC sequence complementary to the sequence of spliceosome RNA U2 served as the branching point sequence. The distance from branching sequence to 3’-SS and the length of polypyrimidine sequence were also chosen as the most frequently found in introns [27]. Restrictase BbsI sequence was also enclosed into intron for further modifications.

The sequence TAAGT(N)_12_YCTAGYNGTCTTC(N)_39_TACTAAC(N)_4_(Y)_15_CAG represents the common sequence of artificial intros. Introns 99 intron N were combined in dimers (in order to increase the number of intron variants) and the obtained library was cloned into plasmid pSB/IR-CA-HybIntr-SV40intr-BleoR-T2A-mCherry. The generated plasmid library was transfected into HEK293T cell line with subsequent selection with zeocin and choice of clones resistant to zeocin and hence containing introns capable of splicing.

From the clones correctly expressing zeocin resistance protein 250 intron variants were extracted for further verification of splicing efficiency and capability to enhance gene expression. The generated introns were sequenced. And as a result of this sequencing it was found that only 12 introns had full-size 99 nucleotide length, and the rest were 48 nucleotides long, but contained 5’-SS and 3’-SS sequences, and polypyrimidine tract. In some introns the branching point sequence TACTAAC was absent.

For the verification of the characteristics of 12 full size introns they were recloned into plasmid pSB-IR-pA-Pause_Site-CAG-TurboGFP-MODC-IntrHBB-bGHpA-SV40pr-BleoR-SV40pA containing fluorescent protein TurboGFP gene (Fig 3b), so that TurboGFP was capable of fluorescence only if correct intron splicing occurred. Capability of the enhancement of protein expression was checked by means of fluorometry; plasmid variants without intron and with SV40 intron were used as controls. Fluorescence levels are shown at Figure 3a. All generated introns were capable to enhance the efficiency of expression 8-12 fold that is comparable with expression enhancement with SV40 intron. Differences in the enhancement induced by individual introns were practically unobserved.

**Figure 3.**
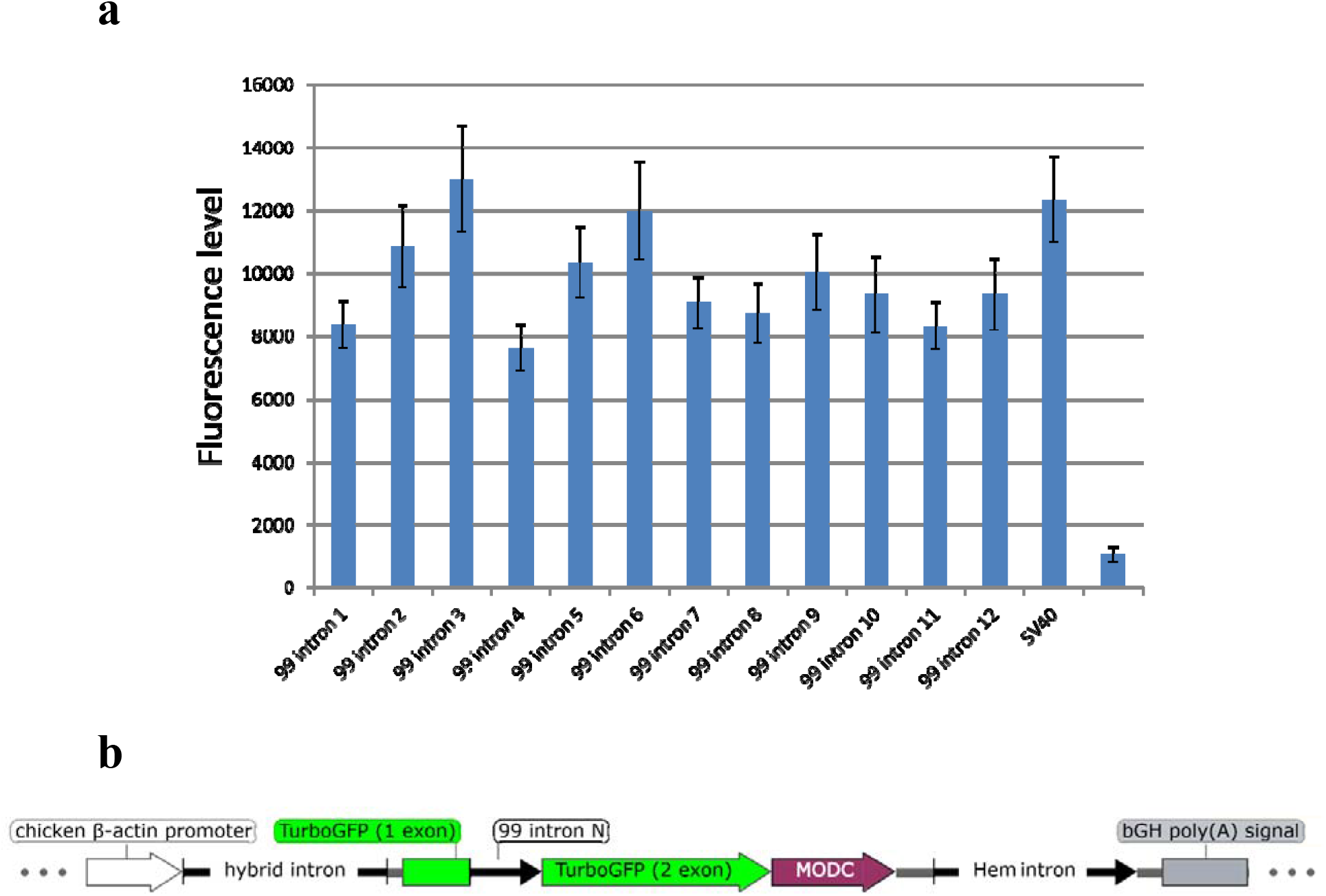
Fluorescence measurement of TurboGFP protein containing the studied introns (a); scheme of the cassette applied for expression investigations (b).

Thus, we have generated a set of introns possessing high splicing efficiency and capable of effective gene expression enhancement.

## Competing interests

The authors declare no competing interests.

## Acknowledgement

The study was supported by RFBR in the frames of scientific project 19-34-60031.

## Notes

### Competing Interest Statement

The authors have declared no competing interest.

