## Supplement for "Artificial introns for effective expression of transgenes in mammalian cells"

pSB/IR-CA-HybIntr-SV40intr-BleoR-T2A-mCherry
Sequence:
AACGCCAGCAACGCGGCCTTTTTACGGTTCCTGGCCTTTTGCTGGCCTTTTGCTCACATGTTCTTTCCTGCGTTATCCCCTGATTCTGTGGATAA
CCGTATTACCGCCTTTGAGTGAGCTGATACCGCTCGCCGCAGCCGAACGACCGAGCGCAGCGAGTCAGTGAGCGAGGAGCCCGATCCCTATA
CAGTTGAAGTCGGAAGTTTACATACACCTTAGCCAAATACATTTAAACTCACTTTTTCACAATTCCTGACATTTAATCCTAGTAAAAATTCCCT
GTCTTAGGTCAGTTAGGATCACCACTTTATTTTAAGAATGTGAAATATCAGAATAATAGTAGAGAGAATGATTCATTTCAGCTTTTATTTCTTT
CATCACATTCCCAGTGGGTCAGAAGTTTACATACACTCAATTAGTATTTGGTAGCATTGCCTTTAAATTGTTTAACTTGGGTCAAACATTTCGA
GTAGCCTTCCACAAGCTAGATCTGGCCATCTAGAGCCCGTTACATAACTTACGGTAAATGGCCCGCCTGGCTGACCGCCCAACGACCCCCGCC
CATTGACGTCAATAGTAACGCCAATAGGGACTTTCCATTGACGTCAATGGGTGGAGTATTTACGGTAAACTGCCCACTTGGCAGTACATCAAG
TGTATCATATGCCAAGTACGCCCCCTATTGACGTCAATGACGGTAAATGGCCCGCCTGGCATTGTGCCCAGTACATGACCTTATGGGACTTTC
CTACTTGGCAGTACATCTACGTATTAGTCATCGCTATTACCATGGTCGAGGTGAGCCCCACGTTCTGCTTCACTCTCCCCATCTCCCCCCCCTC
CCCACCCCCAATTTTGTATTTATTTATTTTTTAATTATTTTGTGCAGCGATGGGGGCGGGGGGGGGGGGGGGGCGCGCGCCAGGCGGGGCGGG
GCGGGGCGAGGGGCGGGGCGGGGCGAGGCGGAGAGGTGCGGCGGCAGCCAATCAGAGCGGCGCGCTCCGAAAGTTTCCTTTTATGGCGAGG
CGGCGGCGGCGGCGGCCCTATAAAAAGCGAAGCGCGCGGCGGGCGGGAGTCGCTGCGACGCTGCCTTCGCCCCGTGCCCCGCTCCGCCGCCG
CCTCGCGCCGCCCGCCCCGGCTCTGACTGACCGCGTTACTCCCACAGGTGAGCGGGCGGGACGGCCCTTCTCCTCCGGGCTGTAATTAGCTGA
GCAAGAGGTAAGGGTTTAAGGGATGGTTGGTTGGTGGGGTATTAATGTTTAATTACCTGGAGCACCTGCCTGAAATCACTTTTTTTCAGGTTT
AACCGGTGCCACCATGGGTACCCCCGGGCCCGGGGAAGACCAGCTGTCAGGTAAGTTTAGTCTTTTTGTCTTTTATTTCAGGTCCCGGATCCG
GTGGTGGTGCAAATCAAAGAACTGCTCCTCAGTTAATGTTGCCTTTACTTCTAGGGAGGCGGGGGTGGAGCCAAGTTGACCAGTGCCGTTCCG
GTGCTCACCGCGCGCGACGTCGCCGGAGCGGTCGAGTTCTGGACCGACCGGCTCGGGTTCTCCCGGGACTTCGTGGAGGACGACTTCGCCGG
TGTGGTCCGGGACGACGTGACCCTGTTCATCAGCGCGGTCCAGGACCAGGTGGTGCCGGACAACACCCTGGCCTGGGTGTGGGTGCGCGGCC
TGGACGAGCTGTACGCCGAGTGGTCGGAGGTCGTGTCCACGAACTTCCGGGACGCCTCCGGGCCGGCCATGACCGAGATCGGCGAGCAGCCG
TGGGGGCGGGAGTTCGCCCTGCGCGACCCGGCCGGCAACTGCGTGCACTTCGTGGCCGAGGAGCAGGACAAGCTTGAGGGCAGAGGAAGTC
TGCTAACATGCGGTGACGTGGAGGAGAATCCCGGCCCTGCTAGCGTGAGCAAGGGCGAGGAGGATAACATGGCTATCATCAAGGAGTTCAT
GCGCTTCAAGGTGCACATGGAGGGCTCCGTGAACGGCCACGAGTTCGAGATCGAGGGCGAGGGCGAGGGCCGCCCCTACGAGGGCACCCAG
ACCGCCAAGCTGAAGGTGACCAAGGGTGGCCCCCTGCCCTTCGCCTGGGACATCCTGTCCCCTCAGTTCATGTACGGCTCCAAGGCCTACGTG
AAGCACCCCGCCGACATCCCCGACTACTTGAAGCTGTCCTTCCCCGAGGGCTTCAAGTGGGAGCGCGTGATGAACTTCGAGGACGGCGGCGT
GGTGACCGTGACCCAGGATTCCTCCTTGCAGGACGGCGAGTTCATCTACAAGGTGAAGCTGCGCGGCACCAACTTCCCCTCCGACGGCCCCG
TAATGCAGAAAAAGACCATGGGCTGGGAGGCCTCCTCCGAGCGGATGTACCCCGAGGACGGCGCCCTGAAGGGCGAGATCAAGCAGAGGCT
GAAGCTGAAGGACGGCGGCCACTACGACGCTGAGGTCAAGACCACCTACAAGGCCAAGAAGCCCGTGCAGGTGCCCGGCGCCTACAACGTC
AACATCAAGTTGGACATCACCTCCCACAACGAGGACTACACCATCGTGGAACAGTACGAACGCGCCGAGGGCCGCCACTCCACCGGCGGCAT
GGACGAGCTGTACAAGGTCGACGGGTAACTAGTGAATTCCATGGATATCAAGCTTCTAAAGCCATGACATCATTTTCTGGAATTTTCCAAGCT
GTTTAAAGGCACAGTCAACTTAGTGTATGTAAACTTCTGACCCACTGGAATTGTGATACAGTGAATTATAAGTGAAATAATCTGTCTGTAAAC
AATTGTTGGAAAAATGACTTGTGTCATGCACAAAGTAGATGTCCTAACTGACTTGCCAAAACTATTGTTTGTTAACAAGAAATTTGTGGAGTA
GTTGAAAAACGAGTTTTAATGACTCCAACTTAAGTGTATGTAAACTTCCGACTTCAACTGTATAGGGATCGGGCACGAAAGGGCCTCGTGATA
CGCCTATTTTTATAGGTTAATGTCATGATAATAATGGTTTCTTAGACGTCAGGTGGCACTTTTCGGGGAAATGTGCGCGGAACCCCTATTTGTT

TATTTTTCTAAATACATTCAAATATGTATCCGCTCATGAGACAATAACCCTGATAAATGCTTCAATAATATTGAAAAAGGAAGAGTATGAGTA
TTCAACATTTCCGTGTCGCCCTTATTCCCTTTTTTGCGGCATTTTGCCTTCCTGTTTTTGCTCACCCAGAAACGCTGGTGAAAGTAAAAGATGCT
GAAGATCAGTTGGGTGCACGAGTGGGTTACATCGAACTGGATCTCAACAGCGGTAAGATCCTTGAGAGTTTTCGCCCCGAAGAACGTTTTCC
AATGATGAGCACTTTTAAAGTTCTGCTATGTGGCGCGGTATTATCCCGTATTGACGCCGGGCAAGAGCAACTCGGTCGCCGCATACACTATTC
TCAGAATGACTTGGTTGAGTACTCACCAGTCACAGAAAAGCATCTTACGGATGGCATGACAGTAAGAGAATTATGCAGTGCTGCCATAACCA
TGAGTGATAACACTGCGGCCAACTTACTTCTGACAACGATCGGAGGACCGAAGGAGCTAACCGCTTTTTTGCACAACATGGGGGATCATGTA
ACTCGCCTTGATCGTTGGGAACCGGAGCTGAATGAAGCCATACCAAACGACGAGCGTGACACCACGATGCCTGTAGCAATGGCAACAACGTT
GCGCAAACTATTAACTGGCGAACTACTTACTCTAGCTTCCCGGCAACAATTAATAGACTGGATGGAGGCGGATAAAGTTGCAGGACCACTTC
TGCGCTCGGCCCTTCCGGCTGGCTGGTTTATTGCTGATAAATCTGGAGCCGGTGAGCGTGGGTCTCGCGGTATCATTGCAGCACTGGGGCCAG
ATGGTAAGCCCTCCCGTATCGTAGTTATCTACACGACGGGGAGTCAGGCAACTATGGATGAACGAAATAGACAGATCGCTGAGATAGGTGCC
TCACTGATTAAGCATTGGTAACTGTCAGACCAAGTTTACTCATATATACTTTAGATTGATTTAAAACTTCATTTTTAATTTAAAAGGATCTAGG
TGAAGATCCTTTTTGATAATCTCATGACCAAAATCCCTTAACGTGAGTTTTCGTTCCACTGAGCGTCAGACCCCGTAGAAAAGATCAAAGGAT
CTTCTTGAGATCCTTTTTTTCTGCGCGTAATCTGCTGCTTGCAAACAAAAAAACCACCGCTACCAGCGGTGGTTTGTTTGCCGGATCAAGAGCT
ACCAACTCTTTTTCCGAAGGTAACTGGCTTCAGCAGAGCGCAGATACCAAATACTGTTCTTCTAGTGTAGCCGTAGTTAGGCCACCACTTCAA
GAACTCTGTAGCACCGCCTACATACCTCGCTCTGCTAATCCTGTTACCAGTGGCTGCTGCCAGTGGCGATAAGTCGTGTCTTACCGGGTTGGA
CTCAAGACGATAGTTACCGGATAAGGCGCAGCGGTCGGGCTGAACGGGGGGTTCGTGCACACAGCCCAGCTTGGAGCGAACGACCTACACC
GAACTGAGATACCTACAGCGTGAGCTATGAGAAAGCGCCACGCTTCCCGAAGGGAGAAAGGCGGACAGGTATCCGGTAAGCGGCAGGGTCG
GAACAGGAGAGCGCACGAGGGAGCTTCCAGGGGGAAACGCCTGGTATCTTTATAGTCCTGTCGGGTTTCGCCACCTCTGACTTGAGCGTCGA
TTTTTGTGATGCTCGTCAGGGGGGCGGAGCCTATGGAAA

pSB/IR-CA-HybIntr-2x99intr-SV40intr-BleoR-T2A-mCherry
Sequence:
AACGCCAGCAACGCGGCCTTTTTACGGTTCCTGGCCTTTTGCTGGCCTTTTGCTCACATGTTCTTTCCTGCGTTATCCCCTGATTCTGTGGATAA
CCGTATTACCGCCTTTGAGTGAGCTGATACCGCTCGCCGCAGCCGAACGACCGAGCGCAGCGAGTCAGTGAGCGAGGAGCCCGATCCCTATA
CAGTTGAAGTCGGAAGTTTACATACACCTTAGCCAAATACATTTAAACTCACTTTTTCACAATTCCTGACATTTAATCCTAGTAAAAATTCCCT
GTCTTAGGTCAGTTAGGATCACCACTTTATTTTAAGAATGTGAAATATCAGAATAATAGTAGAGAGAATGATTCATTTCAGCTTTTATTTCTTT
CATCACATTCCCAGTGGGTCAGAAGTTTACATACACTCAATTAGTATTTGGTAGCATTGCCTTTAAATTGTTTAACTTGGGTCAAACATTTCGA
GTAGCCTTCCACAAGCTAGATCTGGCCATCTAGAGCCCGTTACATAACTTACGGTAAATGGCCCGCCTGGCTGACCGCCCAACGACCCCCGCC
CATTGACGTCAATAGTAACGCCAATAGGGACTTTCCATTGACGTCAATGGGTGGAGTATTTACGGTAAACTGCCCACTTGGCAGTACATCAAG
TGTATCATATGCCAAGTACGCCCCCTATTGACGTCAATGACGGTAAATGGCCCGCCTGGCATTGTGCCCAGTACATGACCTTATGGGACTTTC
CTACTTGGCAGTACATCTACGTATTAGTCATCGCTATTACCATGGTCGAGGTGAGCCCCACGTTCTGCTTCACTCTCCCCATCTCCCCCCCCTC
CCCACCCCCAATTTTGTATTTATTTATTTTTTAATTATTTTGTGCAGCGATGGGGGCGGGGGGGGGGGGGGGGCGCGCGCCAGGCGGGGCGGG
GCGGGGCGAGGGGCGGGGCGGGGCGAGGCGGAGAGGTGCGGCGGCAGCCAATCAGAGCGGCGCGCTCCGAAAGTTTCCTTTTATGGCGAGG
CGGCGGCGGCGGCGGCCCTATAAAAAGCGAAGCGCGCGGCGGGCGGGAGTCGCTGCGACGCTGCCTTCGCCCCGTGCCCCGCTCCGCCGCCG

CCTCGCGCCGCCCGCCCCGGCTCTGACTGACCGCGTTACTCCCACAGGTGAGCGGGCGGGACGGCCCTTCTCCTCCGGGCTGTAATTAGCTGA
GCAAGAGGTAAGGGTTTAAGGGATGGTTGGTTGGTGGGGTATTAATGTTTAATTACCTGGAGCACCTGCCTGAAATCACTTTTTTTCAGGTTT
AACCGGTGCCACCATGGGTACCG
(Introne Sequence)
CTGGCGCGGACCG
(Introne Sequence)
CTGTCAGGTAAGTTTAGTCTTTTTGTCTTTTATTTCAGGTCCCGGATCCGGTGGTGGTGCAAATCAAAGAACTGCTCCTCAGTTAATGTTGCCT
TTACTTCTAGGGAGGCGGGGGTGGAGCCAAGTTGACCAGTGCCGTTCCGGTGCTCACCGCGCGCGACGTCGCCGGAGCGGTCGAGTTCTGGA
CCGACCGGCTCGGGTTCTCCCGGGACTTCGTGGAGGACGACTTCGCCGGTGTGGTCCGGGACGACGTGACCCTGTTCATCAGCGCGGTCCAG
GACCAGGTGGTGCCGGACAACACCCTGGCCTGGGTGTGGGTGCGCGGCCTGGACGAGCTGTACGCCGAGTGGTCGGAGGTCGTGTCCACGAA
CTTCCGGGACGCCTCCGGGCCGGCCNNHACCGAGATCGGCGAGCAGCCGTGGGGGCGGGAGTTCGCCCTGCGCGACCCGGCCGGCAACTGC
GTGCACTTCGTGGCCGAGGAGCAGGACAAGCTTGAGGGCAGAGGAAGTCTGCTAACATGCGGTGACGTGGAGGAGAATCCCGGCCCTGCTA
GCGTGAGCAAGGGCGAGGAGGATAACATGGCTATCATCAAGGAGTTCATGCGCTTCAAGGTGCACATGGAGGGCTCCGTGAACGGCCACGA
GTTCGAGATCGAGGGCGAGGGCGAGGGCCGCCCCTACGAGGGCACCCAGACCGCCAAGCTGAAGGTGACCAAGGGTGGCCCCCTGCCCTTC
GCCTGGGACATCCTGTCCCCTCAGTTCATGTACGGCTCCAAGGCCTACGTGAAGCACCCCGCCGACATCCCCGACTACTTGAAGCTGTCCTTC
CCCGAGGGCTTCAAGTGGGAGCGCGTGATGAACTTCGAGGACGGCGGCGTGGTGACCGTGACCCAGGATTCCTCCTTGCAGGACGGCGAGTT
CATCTACAAGGTGAAGCTGCGCGGCACCAACTTCCCCTCCGACGGCCCCGTAATGCAGAAAAAGACCATGGGCTGGGAGGCCTCCTCCGAGC
GGATGTACCCCGAGGACGGCGCCCTGAAGGGCGAGATCAAGCAGAGGCTGAAGCTGAAGGACGGCGGCCACTACGACGCTGAGGTCAAGAC
CACCTACAAGGCCAAGAAGCCCGTGCAGGTGCCCGGCGCCTACAACGTCAACATCAAGTTGGACATCACCTCCCACAACGAGGACTACACCA
TCGTGGAACAGTACGAACGCGCCGAGGGCCGCCACTCCACCGGCGGCATGGACGAGCTGTACAAGGTCGACGGGTAACTAGTGAATTCCAT
GGATATCAAGCTTCTAAAGCCATGACATCATTTTCTGGAATTTTCCAAGCTGTTTAAAGGCACAGTCAACTTAGTGTATGTAAACTTCTGACCC
ACTGGAATTGTGATACAGTGAATTATAAGTGAAATAATCTGTCTGTAAACAATTGTTGGAAAAATGACTTGTGTCATGCACAAAGTAGATGTC
CTAACTGACTTGCCAAAACTATTGTTTGTTAACAAGAAATTTGTGGAGTAGTTGAAAAACGAGTTTTAATGACTCCAACTTAAGTGTATGTAA
ACTTCCGACTTCAACTGTATAGGGATCGGGCACGAAAGGGCCTCGTGATACGCCTATTTTTATAGGTTAATGTCATGATAATAATGGTTTCTT
AGACGTCAGGTGGCACTTTTCGGGGAAATGTGCGCGGAACCCCTATTTGTTTATTTTTCTAAATACATTCAAATATGTATCCGCTCATGAGAC
AATAACCCTGATAAATGCTTCAATAATATTGAAAAAGGAAGAGTATGAGTATTCAACATTTCCGTGTCGCCCTTATTCCCTTTTTTGCGGCATT
TTGCCTTCCTGTTTTTGCTCACCCAGAAACGCTGGTGAAAGTAAAAGATGCTGAAGATCAGTTGGGTGCACGAGTGGGTTACATCGAACTGGA
TCTCAACAGCGGTAAGATCCTTGAGAGTTTTCGCCCCGAAGAACGTTTTCCAATGATGAGCACTTTTAAAGTTCTGCTATGTGGCGCGGTATT
ATCCCGTATTGACGCCGGGCAAGAGCAACTCGGTCGCCGCATACACTATTCTCAGAATGACTTGGTTGAGTACTCACCAGTCACAGAAAAGC
ATCTTACGGATGGCATGACAGTAAGAGAATTATGCAGTGCTGCCATAACCATGAGTGATAACACTGCGGCCAACTTACTTCTGACAACGATC
GGAGGACCGAAGGAGCTAACCGCTTTTTTGCACAACATGGGGGATCATGTAACTCGCCTTGATCGTTGGGAACCGGAGCTGAATGAAGCCAT
ACCAAACGACGAGCGTGACACCACGATGCCTGTAGCAATGGCAACAACGTTGCGCAAACTATTAACTGGCGAACTACTTACTCTAGCTTCCC
GGCAACAATTAATAGACTGGATGGAGGCGGATAAAGTTGCAGGACCACTTCTGCGCTCGGCCCTTCCGGCTGGCTGGTTTATTGCTGATAAAT
CTGGAGCCGGTGAGCGTGGGTCTCGCGGTATCATTGCAGCACTGGGGCCAGATGGTAAGCCCTCCCGTATCGTAGTTATCTACACGACGGGG
AGTCAGGCAACTATGGATGAACGAAATAGACAGATCGCTGAGATAGGTGCCTCACTGATTAAGCATTGGTAACTGTCAGACCAAGTTTACTC

ATATATACTTTAGATTGATTTAAAACTTCATTTTTAATTTAAAAGGATCTAGGTGAAGATCCTTTTTGATAATCTCATGACCAAAATCCCTTAA
CGTGAGTTTTCGTTCCACTGAGCGTCAGACCCCGTAGAAAAGATCAAAGGATCTTCTTGAGATCCTTTTTTTCTGCGCGTAATCTGCTGCTTGC
AAACAAAAAAACCACCGCTACCAGCGGTGGTTTGTTTGCCGGATCAAGAGCTACCAACTCTTTTTCCGAAGGTAACTGGCTTCAGCAGAGCG
CAGATACCAAATACTGTTCTTCTAGTGTAGCCGTAGTTAGGCCACCACTTCAAGAACTCTGTAGCACCGCCTACATACCTCGCTCTGCTAATCC
TGTTACCAGTGGCTGCTGCCAGTGGCGATAAGTCGTGTCTTACCGGGTTGGACTCAAGACGATAGTTACCGGATAAGGCGCAGCGGTCGGGC
TGAACGGGGGGTTCGTGCACACAGCCCAGCTTGGAGCGAACGACCTACACCGAACTGAGATACCTACAGCGTGAGCTATGAGAAAGCGCCA
CGCTTCCCGAAGGGAGAAAGGCGGACAGGTATCCGGTAAGCGGCAGGGTCGGAACAGGAGAGCGCACGAGGGAGCTTCCAGGGGGAAACG
CCTGGTATCTTTATAGTCCTGTCGGGTTTCGCCACCTCTGACTTGAGCGTCGATTTTTGTGATGCTCGTCAGGGGGGCGGAGCCTATGGAAA

pSB-IR-pA-Pause_Site-CAG-TurboGFP-MODC-IntrHBB-bGHpA-SV40pr-BleoR-SV40pA
Sequence:
CACATTTCCCCGAAAAGTGCCACCTGACGTCTAAGAAACCATTATTATCATGACATTAACCTATAAAAATAGGCGTATCACGAGGCCCTTTCG
TGCCCGATCCCTATACAGTTGAAGTCGGAAGTTTACATACACTTAAGTTGGAGTCATTAAAACTCGTTTTTCAACTACTCCACAAATTTCTTGT
TAACAAACAATAGTTTTGGCAAGTCAGTTAGGACATCTACTTTGTGCATGACACAAGTCATTTTTCCAACAATTGTTTACAGACAGATTATTTC
ACTTATAATTCACTGTATCACAATTCCAGTGGGTCAGAAGTTTACATACACTAAGTTGACTGTGCCTTTAAACAGCTTGGAAAATTCCAGAAA
ATGATGTCATGGCTTTAGCGCGAAGACGCTAGCGAGCTCCCAATAAAATATCTTTATTTTCATTACATCTGTGTGTTGGTTTTTTGTGTGAATC
GATAGTACTAACATACGCTCTCCATCAAAACAAAACGAAACAAAACAAACTAGCAAAATAGGCTGTCCCCAGTGCAAGTGCAGGTGCCAGA
ACATTTCTCTAGCGCTGCTAGCGAAGACAAGATAGACATTGATTATTGACTAGTTATTAATAGTAATCAATTACGGGGTCATTAGTTCATAGC
CCATATATGGAGTTCCGCGTTACATAACTTACGGTAAATGGCCCGCCTGGCTGACCGCCCAACGACCCCCGCCCATTGACGTCAATAATGACG
TATGTTCCCATAGTAACGCCAATAGGGACTTTCCATTGACGTCAATGGGTGGAGTATTTACGGTAAACTGCCCACTTGGCAGTACATCAAGTG
TATCATATGCCAAGTACGCCCCCTATTGACGTCAATGACGGTAAATGGCCCGCCTGGCATTATGCCCAGTACATGACCTTATGGGACTTTCCT
ACTTGGCAGTACATCTACGTATTAGTCATCGCTATTACCATGGTCGAGGTGAGCCCCACGTTCTGCTTCACTCTCCCCATCTCCCCCCCCTCCC
CACCCCCAATTTTGTATTTATTTATTTTTTAATTATTTTGTGCAGCGATGGGGGCGGGGGGGGGGGGGGGGCGCGCGCCAGGCGGGGCGGGGC
GGGGCGAGGGGCGGGGCGGGGCGAGGCGGAGAGGTGCGGCGGCAGCCAATCAGAGCGGCGCGCTCCGAAAGTTTCCTTTTATGGCGAGGCG
GCGGCGGCGGCGGCCCTATAAAAAGCGAAGCGCGCGGCGGGCGGGAGTCGCTGCGACGCTGCCTTCGCCCCGTGCCCCGCTCCGCCGCCGCC
TCGCGCCGCCCGCCCCGGCTCTGACTGACCGCGTTACTCCCACAGGTGAGCGGGCGGGACGGCCCTTCTCCTCCGGGCTGTAATTAGCTGAGC
AAGAGGTAAGGGTTTAAGGGATGGTTGGTTGGTGGGGTATTAATGTTTAATTACCTGGAGCACCTGCCTGAAATCACTTTTTTTCAGGTTGGA
CCGGTCGCCACCATGGAGAGCGACGAGAGCGGCCTGCCCGCCATGGAGATCGAGTGCCGCATCACCGGCACCCTGAACGGCGTGGAGTTCG
AGCTGGTGGGCGGCGGAGAGGGCACCCCCGAGCAGGGCCGCATGACCAACAAGATGAAGAGCACCAAAGGCGCCCTGACCTTCAGCCCCTA
CCTGCTGAGCCACGTGATGGGCTACGGCTTCTACCACTTCGGCACCTACCCCAGCGGCTACGAGAACCCCTTCCTGCACGCCATCAACAACGG
CGGCTACACCAACACCCGCATCGAGAAGTACGAGGACGGCGGCGTGCTGCACGTGAGCTTCAGCTACCGCTACGAGGCCGGCCGCGTGATCG
GCGACTTCAAGGTGATGGGCACCGGCTTCCCCGAGGACAGCGTGATCTTCACCGACAAGATCATCCGCAGCAACGCCACCGTGGAGCACCTG
CACCCCATGGGCGATAACGATCTGGATGGCAGCTTCACCCGCACCTTCAGCCTGCGCGACGGCGGCTACTACAGCTCCGTGGTGGACAGCCA
CATGCACTTCAAGAGCGCCATCCACCCCAGCATCCTGCAGAACGGGGGCCCCATGTTCGCCTTCCGCCGCGTGGAGGAGGATCACAGCAACA

CCGAGCTGGGCATCGTGGAGTACCAGCACGCCTTCAAGACCCCGGATGCAGATGCCGGTGAAGAAAGATCTCGAGATATCAGCCATGGCTTC
CCGCCGGCGGTGGCGGCGCAGGATGATGGCACGCTGCCCATGTCTTGTGCCCAGGAGAGCGGGATGGACCGTCACCCTGCAGCCTGTGCTTC
TGCTAGGATCAATGTGTAGGCGGCCGCGTGACAAGCTGCACGTGGATCCTGAGAACTTCAGGGTGAGTCTATGGGACCCTTGATGTTTTCTTT
CCCCTTCTTTTCTATGGTTAAGTTCATGTCATAGGAAGGGGATAAGTAACAGGGTACAGTTTAGAATGGGAAACAGACGAATGATTGCATCA
GTGTGGAAGTCTCAGGATCGTTTTAGTTTCTTTTATTTGCTGTTCATAACAATTGTTTTCTTTTGTTTAATTCTTGCTTTCTTTTTTTTTCTTCTCC
GCAATTTTTACTATTATACTTAATGCCTTAACATTGTGTATAACAAAAGGAAATATCTCTGAGATACATTAAGTAACTTAAAAAAAAACTTTA
CACAGTCTGCCTAGTACATTACTATTTGGAATATATGTGTGCTTATTTGCATATTCATAATCTCCCTACTTTATTTTCTTTTATTTTTAATTGATA
CATAATCATTATACATATTTATGGGTTAAAGTGTAATGTTTTAATATGTGTACACATATTGACCAAATCAGGGTAATTTTGCATTTGTAATTTT
AAAAAATGCTTTCTTCTTTTAATATACTTTTTTGTTTATCTTATTTCTAATACTTTCCCTAATCTCTTTCTTTCAGGGCAATAATGATACAATGTA
TCATGCCTCTTTGCACCATTCTAAAGAATAACAGTGATAATTTCTGGGTTAAGGCAATAGCAATATTTCTGCATATAAATATTTCTGCATATAA
ATTGTAACTGATGTAAGAGGTTTCATATTGCTAATAGCAGCTACAATCCAGCTACCATTCTGCTTTTATTTTATGGTTGGGATAAGGCTGGATT
ATTCTGAGTCCAAGCTAGGCCCTTTTGCTAATCATGTTCATACTTCTTATCTTCCTCCCACAGCTCCTGGGCAACGTGCTGGTCTGTGTGCTGG
CCCATCACTTTGGCAAAGAATTCACCCCACCAGTGCAGGCTGCCTATCAGAAAGTGGTGGCTGGTGTGGCTAATGCCCTGGCCCACAAGTATC
ACTAAGCTCGCTTTCTTGCTGTCCAATTTCTATTAAAGGTTCCTTTGTTCCCTAAGTCCAACTACTAAACTGGGGGATATTATGAAGGGCCTTG
AGCATTGGATTCTGCCTCTAGATCATAATCAGCCATACCACATTTGTAGAGGTTTTACTTGCTTTAAAAAACCTCCCACACCTCCCCCTGAACC
TGAAACATAAAATGAATGCAATTGTTGTTGTTCTCGCTGATCAGCCTCGACTGTGCCTTCTAGTTGCCAGCCATCTGTTGTTTGCCCCTCCCCC
GTGCCTTCCTTGACCCTGGAAGGTGCCACTCCCACTGTCCTTTCCTAATAAAATGAGGAAATTGCATCGCATTGTCTGAGTAGGTGTCATTCTA
TTCTGGGGGGTGGGGTGGGGCAGGACAGCAAGGGGGAGGATTGGGAAGAGAATAGCAGGCATGCTGGGGACGAGGTCTCTAGAGGCCTGCT
ATGTGTGTCAGTTAGGGTGTGGAAAGTCCCCAGGCTCCCCAGCAGGCAGAAGTATGCAAAGCATGCATCTCAATTAGTCAGCAACCAGGTGT
GGAAAGTCCCCAGGCTCCCCAGCAGGCAGAAGTATGCAAAGCATGCATCTCAATTAGTCAGCAACCATAGTCCCGCCCCTAACTCCGCCCAT
CCCGCCCCTAACTCCGCCCAGTTCCGCCCATTCTCCGCCCCATGGCTGACTAATTTTTTTTATTTATGCAGAGGCCGAGGCCGCCTCTGCCTCT
GAGCTATTCCAGAAGTAGTGAGGAGGCTTTTTTGGAGGCCTAGGCTTTTGCAAAAAGCTCCCGGGAGCTTGTATATCCATTTTCGGATCTGAT
CAGCACGTGTTGACAATTAATCATCGGCATAGTATATCGGCATAGTATAATACGACAAGGTGAGGAACTAAACCATGGCCAAGTTGACCAGT
GCCGTTCCGGTGCTCACCGCGCGCGACGTCGCCGGAGCGGTCGAGTTCTGGACCGACCGGCTCGGGTTCTCCCGGGACTTCGTGGAGGACGA
CTTCGCCGGTGTGGTCCGGGACGACGTGACCCTGTTCATCAGCGCGGTCCAGGACCAGGTGGTGCCGGACAACACCCTGGCCTGGGTGTGGG
TGCGCGGCCTGGACGAGCTGTACGCCGAGTGGTCGGAGGTCGTGTCCACGAACTTCCGGGACGCCTCCGGGCCGGCCATGACCGAGATCGGC
GAGCAGCCGTGGGGGCGGGAGTTCGCCCTGCGCGACCCGGCCGGCAACTGCGTGCACTTCGTGGCCGAGGAGCAGGACTGACACGTGCTAC
GAGATTTCGATTCCACCGCCGCCTTCTATGAAAGGTTGGGCTTCGGAATCGTTTTCCGGGACGCCGGCTGGATGATCCTCCAGCGCGGGGATC
TCATGCTGGAGTTCTTCGCCCACCCCAACTTGTTTATTGCAGCTTATAATGGTTACAAATAAAGCAATAGCATCACAAATTTCACAAATAAAG
CATTTTTTTCACTGCATTCTAGTTGTGGTTTGTCCAAACTCATCAATGTATCTTAGACTAGTGAAGACACGTCTGAGCAGCTTGTGGAAGGCTA
CTCGAAATGTTTGACCCAAGTTAAACAATTTAAAGGCAATGCTACCAAATACTAATTGAGTGTATGTAAACTTCTGACCCACTGGGAATGTGA
TGAAAGAAATAAAAGCTGAAATGAATCATTCTCTCTACTATTATTCTGATATTTCACATTCTTAAAATAAAGTGGTGATCCTAACTGACCTAA
GACAGGGAATTTTTACTAGGATTAAATGTCAGGAATTGTGAAAAAGTGAGTTTAAATGTATTTGGCTAAGGTGTATGTAAACTTCCGACTTCA
ACTGTATAGGGATCGGGCTCCTCGCTCACTGACTCGCTGCGCTCGGTCGTTCGGCTGCGGCGAGCGGTATCAGCTCACTCAAAGGCGGTAATA
CGGTTATCCACAGAATCAGGGGATAACGCAGGAAAGAACATGTGAGCAAAAGGCCAGCAAAAGGCCAGGAACCGTAAAAAGGCCGCGTTG

CTGGCGTTTTTCCATAGGCTCCGCCCCCCTGACGAGCATCACAAAAATCGACGCTCAAGTCAGAGGTGGCGAAACCCGACAGGACTATAAAG
ATACCAGGCGTTTCCCCCTGGAAGCTCCCTCGTGCGCTCTCCTGTTCCGACCCTGCCGCTTACCGGATACCTGTCCGCCTTTCTCCCTTCGGGA
AGCGTGGCGCTTTCTCATAGCTCACGCTGTAGGTATCTCAGTTCGGTGTAGGTCGTTCGCTCCAAGCTGGGCTGTGTGCACGAACCCCCCGTTC
AGCCCGACCGCTGCGCCTTATCCGGTAACTATCGTCTTGAGTCCAACCCGGTAAGACACGACTTATCGCCACTGGCAGCAGCCACTGGTAACA
GGATTAGCAGAGCGAGGTATGTAGGCGGTGCTACAGAGTTCTTGAAGTGGTGGCCTAACTACGGCTACACTAGAAGAACAGTATTTGGTATC
TGCGCTCTGCTGAAGCCAGTTACCTTCGGAAAAAGAGTTGGTAGCTCTTGATCCGGCAAACAAACCACCGCTGGTAGCGGTGGTTTTTTTGTT
TGCAAGCAGCAGATTACGCGCAGAAAAAAAGGATCTCAAGAAGATCCTTTGATCTTTTCTACGGGGTCTGACGCTCAGTGGAACGAAAACTC
ACGTTAAGGGATTTTGGTCATGAGATTATCAAAAAGGATCTTCACCTAGATCCTTTTAAATTAAAAATGAAGTTTTAAATCAATCTAAAGTAT
ATATGAGTAAACTTGGTCTGACAGTTACCAATGCTTAATCAGTGAGGCACCTATCTCAGCGATCTGTCTATTTCGTTCATCCATAGTTGCCTGA
CTCCCCGTCGTGTAGATAACTACGATACGGGAGGGCTTACCATCTGGCCCCAGTGCTGCAATGATACCGCGAGACCCACGCTCACCGGCTCC
AGATTTATCAGCAATAAACCAGCCAGCCGGAAGGGCCGAGCGCAGAAGTGGTCCTGCAACTTTATCCGCCTCCATCCAGTCTATTAATTGTTG
CCGGGAAGCTAGAGTAAGTAGTTCGCCAGTTAATAGTTTGCGCAACGTTGTTGCCATTGCTACAGGCATCGTGGTGTCACGCTCGTCGTTTGG
TATGGCTTCATTCAGCTCCGGTTCCCAACGATCAAGGCGAGTTACATGATCCCCCATGTTGTGCAAAAAAGCGGTTAGCTCCTTCGGTCCTCC
GATCGTTGTCAGAAGTAAGTTGGCCGCAGTGTTATCACTCATGGTTATGGCAGCACTGCATAATTCTCTTACTGTCATGCCATCCGTAAGATG
CTTTTCTGTGACTGGTGAGTACTCAACCAAGTCATTCTGAGAATAGTGTATGCGGCGACCGAGTTGCTCTTGCCCGGCGTCAATACGGGATAA
TACCGCGCCACATAGCAGAACTTTAAAAGTGCTCATCATTGGAAAACGTTCTTCGGGGCGAAAACTCTCAAGGATCTTACCGCTGTTGAGATC
CAGTTCGATGTAACCCACTCGTGCACCCAACTGATCTTCAGCATCTTTTACTTTCACCAGCGTTTCTGGGTGAGCAAAAACAGGAAGGCAAAA
TGCCGCAAAAAAGGGAATAAGGGCGACACGGAAATGTTGAATACTCATACTCTTCCTTTTTCAATATTATTGAAGCATTTATCAGGGTTATTG
TCTCATGAGCGGATACATATTTGAATGTATTTAGAAAAATAAACAAATAGGGGTTCCGCG

pSB-IR-pA-Pause_Site-CAG-TurboGFP-SV40intr-MODC-IntrHBB-bGHpA-SV40pr-BleoR-SV40pA
Sequence:
CACATTTCCCCGAAAAGTGCCACCTGACGTCTAAGAAACCATTATTATCATGACATTAACCTATAAAAATAGGCGTATCACGAGGCCCTTTCG
TGCCCGATCCCTATACAGTTGAAGTCGGAAGTTTACATACACTTAAGTTGGAGTCATTAAAACTCGTTTTTCAACTACTCCACAAATTTCTTGT
TAACAAACAATAGTTTTGGCAAGTCAGTTAGGACATCTACTTTGTGCATGACACAAGTCATTTTTCCAACAATTGTTTACAGACAGATTATTTC
ACTTATAATTCACTGTATCACAATTCCAGTGGGTCAGAAGTTTACATACACTAAGTTGACTGTGCCTTTAAACAGCTTGGAAAATTCCAGAAA
ATGATGTCATGGCTTTAGCGCGAAGACGCTAGCGAGCTCCCAATAAAATATCTTTATTTTCATTACATCTGTGTGTTGGTTTTTTGTGTGAATC
GATAGTACTAACATACGCTCTCCATCAAAACAAAACGAAACAAAACAAACTAGCAAAATAGGCTGTCCCCAGTGCAAGTGCAGGTGCCAGA
ACATTTCTCTAGCGCTGCTAGCGAAGACAAGATAGACATTGATTATTGACTAGTTATTAATAGTAATCAATTACGGGGTCATTAGTTCATAGC
CCATATATGGAGTTCCGCGTTACATAACTTACGGTAAATGGCCCGCCTGGCTGACCGCCCAACGACCCCCGCCCATTGACGTCAATAATGACG
TATGTTCCCATAGTAACGCCAATAGGGACTTTCCATTGACGTCAATGGGTGGAGTATTTACGGTAAACTGCCCACTTGGCAGTACATCAAGTG
TATCATATGCCAAGTACGCCCCCTATTGACGTCAATGACGGTAAATGGCCCGCCTGGCATTATGCCCAGTACATGACCTTATGGGACTTTCCT
ACTTGGCAGTACATCTACGTATTAGTCATCGCTATTACCATGGTCGAGGTGAGCCCCACGTTCTGCTTCACTCTCCCCATCTCCCCCCCCTCCC
CACCCCCAATTTTGTATTTATTTATTTTTTAATTATTTTGTGCAGCGATGGGGGCGGGGGGGGGGGGGGGCGCGCGCCAGGCGGGGCGGGGC

GGGGCGAGGGGCGGGGCGGGGCGAGGCGGAGAGGTGCGGCGGCAGCCAATCAGAGCGGCGCGCTCCGAAAGTTTCCTTTTATGGCGAGGCG
GCGGCGGCGGCGGCCCTATAAAAAGCGAAGCGCGCGGCGGGCGGGAGTCGCTGCGACGCTGCCTTCGCCCCGTGCCCCGCTCCGCCGCCGCC
TCGCGCCGCCCGCCCCGGCTCTGACTGACCGCGTTACTCCCACAGGTGAGCGGGCGGGACGGCCCTTCTCCTCCGGGCTGTAATTAGCTGAGC
AAGAGGTAAGGGTTTAAGGGATGGTTGGTTGGTGGGGTATTAATGTTTAATTACCTGGAGCACCTGCCTGAAATCACTTTTTTTCAGGTTGGA
CCGGTCGCCACCATGGAGAGCGACGAGAGCGGCCTGCCCGCCATGGAGATCGAGTGCCGCATCACCGGCACCCTGAACGGCGTGGAGTTCG
AGCTGGTGGGCGGCGGAGAGGGCACCCCCGAGCAGGGCCGGTAAGTTTAGTCTTTTTGTCTTTTATTTCAGGTCCCGGATCCGGTGGTGGTGC
AAATCAAAGAACTGCTCCTCAGTGGATGTTGCCTTTACTTCTAGCATGACCAACAAGATGAAGAGCACCAAAGGCGCCCTGACCTTCAGCCC
CTACCTGCTGAGCCACGTGATGGGCTACGGCTTCTACCACTTCGGCACCTACCCCAGCGGCTACGAGAACCCCTTCCTGCACGCCATCAACAA
CGGCGGCTACACCAACACCCGCATCGAGAAGTACGAGGACGGCGGCGTGCTGCACGTGAGCTTCAGCTACCGCTACGAGGCCGGCCGCGTG
ATCGGCGACTTCAAGGTGATGGGCACCGGCTTCCCCGAGGACAGCGTGATCTTCACCGACAAGATCATCCGCAGCAACGCCACCGTGGAGCA
CCTGCACCCCATGGGCGATAACGATCTGGATGGCAGCTTCACCCGCACCTTCAGCCTGCGCGACGGCGGCTACTACAGCTCCGTGGTGGACA
GCCACATGCACTTCAAGAGCGCCATCCACCCCAGCATCCTGCAGAACGGGGGCCCCATGTTCGCCTTCCGCCGCGTGGAGGAGGATCACAGC
AACACCGAGCTGGGCATCGTGGAGTACCAGCACGCCTTCAAGACCCCGGATGCAGATGCCGGTGAAGAAAGATCTCGAGATATCAGCCATG
GCTTCCCGCCGGCGGTGGCGGCGCAGGATGATGGCACGCTGCCCATGTCTTGTGCCCAGGAGAGCGGGATGGACCGTCACCCTGCAGCCTGT
GCTTCTGCTAGGATCAATGTGTAGGCGGCCGCGTGACAAGCTGCACGTGGATCCTGAGAACTTCAGGGTGAGTCTATGGGACCCTTGATGTTT
TCTTTCCCCTTCTTTTCTATGGTTAAGTTCATGTCATAGGAAGGGGATAAGTAACAGGGTACAGTTTAGAATGGGAAACAGACGAATGATTGC
ATCAGTGTGGAAGTCTCAGGATCGTTTTAGTTTCTTTTATTTGCTGTTCATAACAATTGTTTTCTTTTGTTTAATTCTTGCTTTCTTTTTTTTTCTT
CTCCGCAATTTTTACTATTATACTTAATGCCTTAACATTGTGTATAACAAAAGGAAATATCTCTGAGATACATTAAGTAACTTAAAAAAAAAC
TTTACACAGTCTGCCTAGTACATTACTATTTGGAATATATGTGTGCTTATTTGCATATTCATAATCTCCCTACTTTATTTTCTTTTATTTTTAATT
GATACATAATCATTATACATATTTATGGGTTAAAGTGTAATGTTTTAATATGTGTACACATATTGACCAAATCAGGGTAATTTTGCATTTGTAA
TTTTAAAAAATGCTTTCTTCTTTTAATATACTTTTTTGTTTATCTTATTTCTAATACTTTCCCTAATCTCTTTCTTTCAGGGCAATAATGATACAA
TGTATCATGCCTCTTTGCACCATTCTAAAGAATAACAGTGATAATTTCTGGGTTAAGGCAATAGCAATATTTCTGCATATAAATATTTCTGCAT
ATAAATTGTAACTGATGTAAGAGGTTTCATATTGCTAATAGCAGCTACAATCCAGCTACCATTCTGCTTTTATTTTATGGTTGGGATAAGGCTG
GATTATTCTGAGTCCAAGCTAGGCCCTTTTGCTAATCATGTTCATACTTCTTATCTTCCTCCCACAGCTCCTGGGCAACGTGCTGGTCTGTGTGC
TGGCCCATCACTTTGGCAAAGAATTCACCCCACCAGTGCAGGCTGCCTATCAGAAAGTGGTGGCTGGTGTGGCTAATGCCCTGGCCCACAAGT
ATCACTAAGCTCGCTTTCTTGCTGTCCAATTTCTATTAAAGGTTCCTTTGTTCCCTAAGTCCAACTACTAAACTGGGGGATATTATGAAGGGCC
TTGAGCATTGGATTCTGCCTCTAGATCATAATCAGCCATACCACATTTGTAGAGGTTTTACTTGCTTTAAAAAACCTCCCACACCTCCCCCTGA
ACCTGAAACATAAAATGAATGCAATTGTTGTTGTTCTCGCTGATCAGCCTCGACTGTGCCTTCTAGTTGCCAGCCATCTGTTGTTTGCCCCTCC
CCCGTGCCTTCCTTGACCCTGGAAGGTGCCACTCCCACTGTCCTTTCCTAATAAAATGAGGAAATTGCATCGCATTGTCTGAGTAGGTGTCATT
CTATTCTGGGGGGTGGGGTGGGGCAGGACAGCAAGGGGGAGGATTGGGAAGAGAATAGCAGGCATGCTGGGGACGAGGTCTCTAGAGGCCT
GCTATGTGTGTCAGTTAGGGTGTGGAAAGTCCCCAGGCTCCCCAGCAGGCAGAAGTATGCAAAGCATGCATCTCAATTAGTCAGCAACCAGG
TGTGGAAAGTCCCCAGGCTCCCCAGCAGGCAGAAGTATGCAAAGCATGCATCTCAATTAGTCAGCAACCATAGTCCCGCCCCTAACTCCGCC
CATCCCGCCCCTAACTCCGCCCAGTTCCGCCCATTCTCCGCCCCATGGCTGACTAATTTTTTTTATTTATGCAGAGGCCGAGGCCGCCTCTGCC
TCTGAGCTATTCCAGAAGTAGTGAGGAGGCTTTTTTGGAGGCCTAGGCTTTTGCAAAAAGCTCCCGGGAGCTTGTATATCCATTTTCGGATCT
GATCAGCACGTGTTGACAATTAATCATCGGCATAGTATATCGGCATAGTATAATACGACAAGGTGAGGAACTAAACCATGGCCAAGTTGACC

AGTGCCGTTCCGGTGCTCACCGCGCGCGACGTCGCCGGAGCGGTCGAGTTCTGGACCGACCGGCTCGGGTTCTCCCGGGACTTCGTGGAGGA
CGACTTCGCCGGTGTGGTCCGGGACGACGTGACCCTGTTCATCAGCGCGGTCCAGGACCAGGTGGTGCCGGACAACACCCTGGCCTGGGTGT
GGGTGCGCGGCCTGGACGAGCTGTACGCCGAGTGGTCGGAGGTCGTGTCCACGAACTTCCGGGACGCCTCCGGGCCGGCCATGACCGAGATC
GGCGAGCAGCCGTGGGGGCGGGAGTTCGCCCTGCGCGACCCGGCCGGCAACTGCGTGCACTTCGTGGCCGAGGAGCAGGACTGACACGTGC
TACGAGATTTCGATTCCACCGCCGCCTTCTATGAAAGGTTGGGCTTCGGAATCGTTTTCCGGGACGCCGGCTGGATGATCCTCCAGCGCGGGG
ATCTCATGCTGGAGTTCTTCGCCCACCCCAACTTGTTTATTGCAGCTTATAATGGTTACAAATAAAGCAATAGCATCACAAATTTCACAAATA
AAGCATTTTTTTCACTGCATTCTAGTTGTGGTTTGTCCAAACTCATCAATGTATCTTAGACTAGTGAAGACACGTCTGAGCAGCTTGTGGAAGG
CTACTCGAAATGTTTGACCCAAGTTAAACAATTTAAAGGCAATGCTACCAAATACTAATTGAGTGTATGTAAACTTCTGACCCACTGGGAATG
TGATGAAAGAAATAAAAGCTGAAATGAATCATTCTCTCTACTATTATTCTGATATTTCACATTCTTAAAATAAAGTGGTGATCCTAACTGACC
TAAGACAGGGAATTTTTACTAGGATTAAATGTCAGGAATTGTGAAAAAGTGAGTTTAAATGTATTTGGCTAAGGTGTATGTAAACTTCCGACT
TCAACTGTATAGGGATCGGGCTCCTCGCTCACTGACTCGCTGCGCTCGGTCGTTCGGCTGCGGCGAGCGGTATCAGCTCACTCAAAGGCGGTA
ATACGGTTATCCACAGAATCAGGGGATAACGCAGGAAAGAACATGTGAGCAAAAGGCCAGCAAAAGGCCAGGAACCGTAAAAAGGCCGCG
TTGCTGGCGTTTTTCCATAGGCTCCGCCCCCCTGACGAGCATCACAAAAATCGACGCTCAAGTCAGAGGTGGCGAAACCCGACAGGACTATA
AAGATACCAGGCGTTTCCCCCTGGAAGCTCCCTCGTGCGCTCTCCTGTTCCGACCCTGCCGCTTACCGGATACCTGTCCGCCTTTCTCCCTTCG
GGAAGCGTGGCGCTTTCTCATAGCTCACGCTGTAGGTATCTCAGTTCGGTGTAGGTCGTTCGCTCCAAGCTGGGCTGTGTGCACGAACCCCCC
GTTCAGCCCGACCGCTGCGCCTTATCCGGTAACTATCGTCTTGAGTCCAACCCGGTAAGACACGACTTATCGCCACTGGCAGCAGCCACTGGT
AACAGGATTAGCAGAGCGAGGTATGTAGGCGGTGCTACAGAGTTCTTGAAGTGGTGGCCTAACTACGGCTACACTAGAAGAACAGTATTTGG
TATCTGCGCTCTGCTGAAGCCAGTTACCTTCGGAAAAAGAGTTGGTAGCTCTTGATCCGGCAAACAAACCACCGCTGGTAGCGGTGGTTTTTT
TGTTTGCAAGCAGCAGATTACGCGCAGAAAAAAAGGATCTCAAGAAGATCCTTTGATCTTTTCTACGGGGTCTGACGCTCAGTGGAACGAAA
ACTCACGTTAAGGGATTTTGGTCATGAGATTATCAAAAAGGATCTTCACCTAGATCCTTTTAAATTAAAAATGAAGTTTTAAATCAATCTAAA
GTATATATGAGTAAACTTGGTCTGACAGTTACCAATGCTTAATCAGTGAGGCACCTATCTCAGCGATCTGTCTATTTCGTTCATCCATAGTTGC
CTGACTCCCCGTCGTGTAGATAACTACGATACGGGAGGGCTTACCATCTGGCCCCAGTGCTGCAATGATACCGCGAGACCCACGCTCACCGG
CTCCAGATTTATCAGCAATAAACCAGCCAGCCGGAAGGGCCGAGCGCAGAAGTGGTCCTGCAACTTTATCCGCCTCCATCCAGTCTATTAATT
GTTGCCGGGAAGCTAGAGTAAGTAGTTCGCCAGTTAATAGTTTGCGCAACGTTGTTGCCATTGCTACAGGCATCGTGGTGTCACGCTCGTCGT
TTGGTATGGCTTCATTCAGCTCCGGTTCCCAACGATCAAGGCGAGTTACATGATCCCCCATGTTGTGCAAAAAAGCGGTTAGCTCCTTCGGTC
CTCCGATCGTTGTCAGAAGTAAGTTGGCCGCAGTGTTATCACTCATGGTTATGGCAGCACTGCATAATTCTCTTACTGTCATGCCATCCGTAAG
ATGCTTTTCTGTGACTGGTGAGTACTCAACCAAGTCATTCTGAGAATAGTGTATGCGGCGACCGAGTTGCTCTTGCCCGGCGTCAATACGGGA
TAATACCGCGCCACATAGCAGAACTTTAAAAGTGCTCATCATTGGAAAACGTTCTTCGGGGCGAAAACTCTCAAGGATCTTACCGCTGTTGAG
ATCCAGTTCGATGTAACCCACTCGTGCACCCAACTGATCTTCAGCATCTTTTACTTTCACCAGCGTTTCTGGGTGAGCAAAAACAGGAAGGCA
AAATGCCGCAAAAAAGGGAATAAGGGCGACACGGAAATGTTGAATACTCATACTCTTCCTTTTTCAATATTATTGAAGCATTTATCAGGGTTA
TTGTCTCATGAGCGGATACATATTTGAATGTATTTAGAAAAATAAACAAATAGGGGTTCCGCG

pSB-IR-pA-Pause_Site-CAG-TurboGFP-99intr-MODC-IntrHBB-bGHpA-SV40pr-BleoR-SV40pA
Sequence:

CACATTTCCCCGAAAAGTGCCACCTGACGTCTAAGAAACCATTATTATCATGACATTAACCTATAAAAATAGGCGTATCACGAGGCCCTTTCG
TGCCCGATCCCTATACAGTTGAAGTCGGAAGTTTACATACACTTAAGTTGGAGTCATTAAAACTCGTTTTTCAACTACTCCACAAATTTCTTGT
TAACAAACAATAGTTTTGGCAAGTCAGTTAGGACATCTACTTTGTGCATGACACAAGTCATTTTTCCAACAATTGTTTACAGACAGATTATTTC
ACTTATAATTCACTGTATCACAATTCCAGTGGGTCAGAAGTTTACATACACTAAGTTGACTGTGCCTTTAAACAGCTTGGAAAATTCCAGAAA
ATGATGTCATGGCTTTAGCGCGAAGACGCTAGCGAGCTCCCAATAAAATATCTTTATTTTCATTACATCTGTGTGTTGGTTTTTTGTGTGAATC
GATAGTACTAACATACGCTCTCCATCAAAACAAAACGAAACAAAACAAACTAGCAAAATAGGCTGTCCCCAGTGCAAGTGCAGGTGCCAGA
ACATTTCTCTAGCGCTGCTAGCGAAGACAAGATAGACATTGATTATTGACTAGTTATTAATAGTAATCAATTACGGGGTCATTAGTTCATAGC
CCATATATGGAGTTCCGCGTTACATAACTTACGGTAAATGGCCCGCCTGGCTGACCGCCCAACGACCCCCGCCCATTGACGTCAATAATGACG
TATGTTCCCATAGTAACGCCAATAGGGACTTTCCATTGACGTCAATGGGTGGAGTATTTACGGTAAACTGCCCACTTGGCAGTACATCAAGTG
TATCATATGCCAAGTACGCCCCCTATTGACGTCAATGACGGTAAATGGCCCGCCTGGCATTATGCCCAGTACATGACCTTATGGGACTTTCCT
ACTTGGCAGTACATCTACGTATTAGTCATCGCTATTACCATGGTCGAGGTGAGCCCCACGTTCTGCTTCACTCTCCCCATCTCCCCCCCCTCCC
CACCCCCAATTTTGTATTTATTTATTTTTTAATTATTTTGTGCAGCGATGGGGGCGGGGGGGGGGGGGGGGCGCGCGCCAGGCGGGGCGGGGC
GGGGCGAGGGGCGGGGCGGGGCGAGGCGGAGAGGTGCGGCGGCAGCCAATCAGAGCGGCGCGCTCCGAAAGTTTCCTTTTATGGCGAGGCG
GCGGCGGCGGCGGCCCTATAAAAAGCGAAGCGCGCGGCGGGCGGGAGTCGCTGCGACGCTGCCTTCGCCCCGTGCCCCGCTCCGCCGCCGCC
TCGCGCCGCCCGCCCCGGCTCTGACTGACCGCGTTACTCCCACAGGTGAGCGGGCGGGACGGCCCTTCTCCTCCGGGCTGTAATTAGCTGAGC
AAGAGGTAAGGGTTTAAGGGATGGTTGGTTGGTGGGGTATTAATGTTTAATTACCTGGAGCACCTGCCTGAAATCACTTTTTTTCAGGTTGGA
CCGGTCGCCACCATGGAGAGCGACGAGAGCGGCCTGCCCGCCATGGAGATCGAGTGCCGCATCACCGGCACCCTGAACGGCGTGGAGTTCG
AGCTGGTGGGCGGCGGAGAGGGCACCCCCGAGCAGGGCCG
(Introne Sequence)
CATGACCAACAAGATGAAGAGCACCAAAGGCGCCCTGACCTTCAGCCCCTACCTGCTGAGCCACGTGATGGGCTACGGCTTCTACCACTTCG
GCACCTACCCCAGCGGCTACGAGAACCCCTTCCTGCACGCCATCAACAACGGCGGCTACACCAACACCCGCATCGAGAAGTACGAGGACGGC
GGCGTGCTGCACGTGAGCTTCAGCTACCGCTACGAGGCCGGCCGCGTGATCGGCGACTTCAAGGTGATGGGCACCGGCTTCCCCGAGGACAG
CGTGATCTTCACCGACAAGATCATCCGCAGCAACGCCACCGTGGAGCACCTGCACCCCATGGGCGATAACGATCTGGATGGCAGCTTCACCC
GCACCTTCAGCCTGCGCGACGGCGGCTACTACAGCTCCGTGGTGGACAGCCACATGCACTTCAAGAGCGCCATCCACCCCAGCATCCTGCAG
AACGGGGGCCCCATGTTCGCCTTCCGCCGCGTGGAGGAGGATCACAGCAACACCGAGCTGGGCATCGTGGAGTACCAGCACGCCTTCAAGAC
CCCGGATGCAGATGCCGGTGAAGAAAGATCTCGAGATATCAGCCATGGCTTCCCGCCGGCGGTGGCGGCGCAGGATGATGGCACGCTGCCCA
TGTCTTGTGCCCAGGAGAGCGGGATGGACCGTCACCCTGCAGCCTGTGCTTCTGCTAGGATCAATGTGTAGGCGGCCGCGTGACAAGCTGCA
CGTGGATCCTGAGAACTTCAGGGTGAGTCTATGGGACCCTTGATGTTTTCTTTCCCCTTCTTTTCTATGGTTAAGTTCATGTCATAGGAAGGGG
ATAAGTAACAGGGTACAGTTTAGAATGGGAAACAGACGAATGATTGCATCAGTGTGGAAGTCTCAGGATCGTTTTAGTTTCTTTTATTTGCTG
TTCATAACAATTGTTTTCTTTTGTTTAATTCTTGCTTTCTTTTTTTTTCTTCTCCGCAATTTTTACTATTATACTTAATGCCTTAACATTGTGTATA
ACAAAAGGAAATATCTCTGAGATACATTAAGTAACTTAAAAAAAAACTTTACACAGTCTGCCTAGTACATTACTATTTGGAATATATGTGTGC
TTATTTGCATATTCATAATCTCCCTACTTTATTTTCTTTTATTTTTAATTGATACATAATCATTATACATATTTATGGGTTAAAGTGTAATGTTTT
AATATGTGTACACATATTGACCAAATCAGGGTAATTTTGCATTTGTAATTTTAAAAAATGCTTTCTTCTTTTAATATACTTTTTTGTTTATCTTA
TTTCTAATACTTTCCCTAATCTCTTTCTTTCAGGGCAATAATGATACAATGTATCATGCCTCTTTGCACCATTCTAAAGAATAACAGTGATAAT
TTCTGGGTTAAGGCAATAGCAATATTTCTGCATATAAATATTTCTGCATATAAATTGTAACTGATGTAAGAGGTTTCATATTGCTAATAGCAGC

TACAATCCAGCTACCATTCTGCTTTTATTTTATGGTTGGGATAAGGCTGGATTATTCTGAGTCCAAGCTAGGCCCTTTTGCTAATCATGTTCAT
ACTTCTTATCTTCCTCCCACAGCTCCTGGGCAACGTGCTGGTCTGTGTGCTGGCCCATCACTTTGGCAAAGAATTCACCCCACCAGTGCAGGCT
GCCTATCAGAAAGTGGTGGCTGGTGTGGCTAATGCCCTGGCCCACAAGTATCACTAAGCTCGCTTTCTTGCTGTCCAATTTCTATTAAAGGTTC
CTTTGTTCCCTAAGTCCAACTACTAAACTGGGGGATATTATGAAGGGCCTTGAGCATTGGATTCTGCCTCTAGATCATAATCAGCCATACCAC
ATTTGTAGAGGTTTTACTTGCTTTAAAAAACCTCCCACACCTCCCCCTGAACCTGAAACATAAAATGAATGCAATTGTTGTTGTTCTCGCTGAT
CAGCCTCGACTGTGCCTTCTAGTTGCCAGCCATCTGTTGTTTGCCCCTCCCCCGTGCCTTCCTTGACCCTGGAAGGTGCCACTCCCACTGTCCTT
TCCTAATAAAATGAGGAAATTGCATCGCATTGTCTGAGTAGGTGTCATTCTATTCTGGGGGGTGGGGTGGGGCAGGACAGCAAGGGGGAGGA
TTGGGAAGAGAATAGCAGGCATGCTGGGGACGAGGTCTCTAGAGGCCTGCTATGTGTGTCAGTTAGGGTGTGGAAAGTCCCCAGGCTCCCCA
GCAGGCAGAAGTATGCAAAGCATGCATCTCAATTAGTCAGCAACCAGGTGTGGAAAGTCCCCAGGCTCCCCAGCAGGCAGAAGTATGCAAA
GCATGCATCTCAATTAGTCAGCAACCATAGTCCCGCCCCTAACTCCGCCCATCCCGCCCCTAACTCCGCCCAGTTCCGCCCATTCTCCGCCCCA
TGGCTGACTAATTTTTTTTATTTATGCAGAGGCCGAGGCCGCCTCTGCCTCTGAGCTATTCCAGAAGTAGTGAGGAGGCTTTTTTGGAGGCCTA
GGCTTTTGCAAAAAGCTCCCGGGAGCTTGTATATCCATTTTCGGATCTGATCAGCACGTGTTGACAATTAATCATCGGCATAGTATATCGGCA
TAGTATAATACGACAAGGTGAGGAACTAAACCATGGCCAAGTTGACCAGTGCCGTTCCGGTGCTCACCGCGCGCGACGTCGCCGGAGCGGTC
GAGTTCTGGACCGACCGGCTCGGGTTCTCCCGGGACTTCGTGGAGGACGACTTCGCCGGTGTGGTCCGGGACGACGTGACCCTGTTCATCAGC
GCGGTCCAGGACCAGGTGGTGCCGGACAACACCCTGGCCTGGGTGTGGGTGCGCGGCCTGGACGAGCTGTACGCCGAGTGGTCGGAGGTCGT
GTCCACGAACTTCCGGGACGCCTCCGGGCCGGCCATGACCGAGATCGGCGAGCAGCCGTGGGGGCGGGAGTTCGCCCTGCGCGACCCGGCC
GGCAACTGCGTGCACTTCGTGGCCGAGGAGCAGGACTGACACGTGCTACGAGATTTCGATTCCACCGCCGCCTTCTATGAAAGGTTGGGCTTC
GGAATCGTTTTCCGGGACGCCGGCTGGATGATCCTCCAGCGCGGGGATCTCATGCTGGAGTTCTTCGCCCACCCCAACTTGTTTATTGCAGCTT
ATAATGGTTACAAATAAAGCAATAGCATCACAAATTTCACAAATAAAGCATTTTTTTCACTGCATTCTAGTTGTGGTTTGTCCAAACTCATCA
ATGTATCTTAGACTAGTGAAGACACGTCTGAGCAGCTTGTGGAAGGCTACTCGAAATGTTTGACCCAAGTTAAACAATTTAAAGGCAATGCT
ACCAAATACTAATTGAGTGTATGTAAACTTCTGACCCACTGGGAATGTGATGAAAGAAATAAAAGCTGAAATGAATCATTCTCTCTACTATTA
TTCTGATATTTCACATTCTTAAAATAAAGTGGTGATCCTAACTGACCTAAGACAGGGAATTTTTACTAGGATTAAATGTCAGGAATTGTGAAA
AAGTGAGTTTAAATGTATTTGGCTAAGGTGTATGTAAACTTCCGACTTCAACTGTATAGGGATCGGGCTCCTCGCTCACTGACTCGCTGCGCT
CGGTCGTTCGGCTGCGGCGAGCGGTATCAGCTCACTCAAAGGCGGTAATACGGTTATCCACAGAATCAGGGGATAACGCAGGAAAGAACAT
GTGAGCAAAAGGCCAGCAAAAGGCCAGGAACCGTAAAAAGGCCGCGTTGCTGGCGTTTTTCCATAGGCTCCGCCCCCCTGACGAGCATCACA
AAAATCGACGCTCAAGTCAGAGGTGGCGAAACCCGACAGGACTATAAAGATACCAGGCGTTTCCCCCTGGAAGCTCCCTCGTGCGCTCTCCT
GTTCCGACCCTGCCGCTTACCGGATACCTGTCCGCCTTTCTCCCTTCGGGAAGCGTGGCGCTTTCTCATAGCTCACGCTGTAGGTATCTCAGTT
CGGTGTAGGTCGTTCGCTCCAAGCTGGGCTGTGTGCACGAACCCCCCGTTCAGCCCGACCGCTGCGCCTTATCCGGTAACTATCGTCTTGAGT
CCAACCCGGTAAGACACGACTTATCGCCACTGGCAGCAGCCACTGGTAACAGGATTAGCAGAGCGAGGTATGTAGGCGGTGCTACAGAGTTC
TTGAAGTGGTGGCCTAACTACGGCTACACTAGAAGAACAGTATTTGGTATCTGCGCTCTGCTGAAGCCAGTTACCTTCGGAAAAAGAGTTGGT
AGCTCTTGATCCGGCAAACAAACCACCGCTGGTAGCGGTGGTTTTTTTGTTTGCAAGCAGCAGATTACGCGCAGAAAAAAAGGATCTCAAGA
AGATCCTTTGATCTTTTCTACGGGGTCTGACGCTCAGTGGAACGAAAACTCACGTTAAGGGATTTTGGTCATGAGATTATCAAAAAGGATCTT
CACCTAGATCCTTTTAAATTAAAAATGAAGTTTTAAATCAATCTAAAGTATATATGAGTAAACTTGGTCTGACAGTTACCAATGCTTAATCAG
TGAGGCACCTATCTCAGCGATCTGTCTATTTCGTTCATCCATAGTTGCCTGACTCCCCGTCGTGTAGATAACTACGATACGGGAGGGCTTACCA
TCTGGCCCCAGTGCTGCAATGATACCGCGAGACCCACGCTCACCGGCTCCAGATTTATCAGCAATAAACCAGCCAGCCGGAAGGGCCGAGCG

CAGAAGTGGTCCTGCAACTTTATCCGCCTCCATCCAGTCTATTAATTGTTGCCGGGAAGCTAGAGTAAGTAGTTCGCCAGTTAATAGTTTGCG
CAACGTTGTTGCCATTGCTACAGGCATCGTGGTGTCACGCTCGTCGTTTGGTATGGCTTCATTCAGCTCCGGTTCCCAACGATCAAGGCGAGTT
ACATGATCCCCCATGTTGTGCAAAAAAGCGGTTAGCTCCTTCGGTCCTCCGATCGTTGTCAGAAGTAAGTTGGCCGCAGTGTTATCACTCATG
GTTATGGCAGCACTGCATAATTCTCTTACTGTCATGCCATCCGTAAGATGCTTTTCTGTGACTGGTGAGTACTCAACCAAGTCATTCTGAGAAT
AGTGTATGCGGCGACCGAGTTGCTCTTGCCCGGCGTCAATACGGGATAATACCGCGCCACATAGCAGAACTTTAAAAGTGCTCATCATTGGA
AAACGTTCTTCGGGGCGAAAACTCTCAAGGATCTTACCGCTGTTGAGATCCAGTTCGATGTAACCCACTCGTGCACCCAACTGATCTTCAGCA
TCTTTTACTTTCACCAGCGTTTCTGGGTGAGCAAAAACAGGAAGGCAAAATGCCGCAAAAAAGGGAATAAGGGCGACACGGAAATGTTGAAT
ACTCATACTCTTCCTTTTTCAATATTATTGAAGCATTTATCAGGGTTATTGTCTCATGAGCGGATACATATTTGAATGTATTTAGAAAAATAAA
CAAATAGGGGTTCCGCG

Introne 99 1-12
Sequence:
gtaagtgagtcatatactcctagcagtcttcaatgtgaacgactttggaacaagcgttcgcacggcacattactaaccaggctctttcctcttcttcag
gtaagtggatcaacagcatctagttgtcttctgcaaccatcactccttactgaactcacgctcgcggctctactaacgcgaccttccttctccctccag
gtaagtatcactcaactatctagccgtcttcgggacgtcatcacatccagtcaaccgtgggctacgcatctactaacagcgtccttccttctttctcag
gtaagtccccgtactgcccctagccgtcttcttgactcactagcccgacgacagaaacggtactggacggtactaaccgggttttcttttcctcttcag
gtaagttgtgccgtcggccctagtggtcttcgctgcgcaggacaccatgtgtgcacctgccaaggctcgatactaacaatgtctcttttctttccccag
gtaagttggtcagtccgccctagtagtcttctgctgcatgcctgcacggccaaacctgtcacgctgctgctactaacgcagtctcccttccttctccag
gtaagtataactgactggtctagttgtcttcgtatcatcgtggtgttcatggttttgacgctccgatccgtactaacaagacctttccttctccctcag
gtaagtggtgctgcacctcctagcggtcttctgtcagtcggcacgttgccgcaactcacgacttcgcaaatactaaccggcctctcctttctccttcag
gtaagtgtgcagccacgttctagttgtcttctgccacgtccacggctcgcccgtacgcactcgcgcggattactaacatcgtctccttctcttcctcag
gtaagttgttgcgtacagtctagtagtcttcgtgcactgaccatgcctgcaactgccggctggcctagcctactaacatgcctttctccctttctccag
gtaagtgttgggtactcacctagcagtcttcgtcgcacctgcaatgcctgccgggatccgccctcgatcgtactaactggatccttctcccctcttcag
gtaagtagacctgactggcctagccgtcttcgttcacacgtggtccacatggtccctttgactgcactaatactaacgcgttcttcttccttttccag
